# Single adeno-associated virus-based multiplexed CRISPR-Cas9 system to nullify core components of the mammalian molecular clock

**DOI:** 10.1101/2020.07.02.184119

**Authors:** Boil Kim, Jihoon Kim, Minjeong Chun, Inah Park, Mijung Choi, Kyungjin Kim, Han Kyoung Choe

## Abstract

The mammalian molecular clock is based on a transcription-translation feedback loop (TTFL) containing Period1, 2 (*Per1, 2*), Cryptochrome1, 2 (*Cry1, 2*), and Brain and Muscle ARNT-Like 1 (*Bmal1*). TTFL robustness is endowed by genetic complementation between these components; therefore, multiple genes must be knocked out to physiologically investigate the molecular clock, which requires extensive research resources. To facilitate molecular clock disruption, we developed a CRISPR-Cas9-based single adeno-associated viral (AAV) system targeting the circadian clock (CSAC) for *Pers, Crys*, or *Bmal1*. First, we designed single guide RNAs (sgRNAs) targeting individual clock genes using an *in silico* approach and validated their efficiency in Neuro2a cells. To target multiple genes, multiplex sgRNA plasmids were constructed using Golden Gate assembly and expressed in viral vectors. CSAC efficiency was demonstrated by decreased protein expression *in vitro* and ablated molecular oscillation *ex vivo*. We also measured locomotor activity and body temperature in Cas9-expressing mice injected with CSAC at the suprachiasmatic nucleus. Circadian rhythm disruption was observed under free-running conditions, indicating that CSAC can efficiently and robustly disrupt molecular circadian clock. Thus, CSAC is a simple and powerful tool for investigating the physiological role of the molecular clock *in vivo*.

## INTRODUCTION

Circadian rhythm regulates the majority of physiological processes in an organism. In mammals, this system has a hierarchical structure with a master clock machinery located in the suprachiasmatic nucleus (SCN) that synchronizes local clocks in almost every organ, tissue, and cell via humoral and neural signaling^[1,2]^. All individual cellular circadian oscillators, whether they are master or local, are driven by molecular clockwork that operate via transcription-translation negative feedback loops. These loops mainly consist of the positive regulators *Bmal1* and *Clock* and the negative regulators Cryptochrome (*Cry*) and Period (*Per*), which are finely modulated by post-transcriptional and −translational modifications^[3,4]^. Defects in local molecular oscillators can cause the abnormal functioning of the corresponding tissues or organs, resulting in sleep disorders, mood disorders, obesity, and autoimmunity^[5]^. Recent advances in cell categorization based on single-cell RNA sequencing have refined our understanding of cellular organization in the body; however, further studies are required to investigate the role of local circadian oscillators in diverse unappreciated cell types^[6,7]^. The expanding repertoire of *Cre* driver lines has served as a genetic handle for various cell types^[8,9]^; however, an efficient and approachable technique to abolish local clocks in specific cell types is also required to facilitate our understanding of the role of local clocks.

Genetically engineered mice have provided a model system to address the roles of local molecular clocks and the mechanisms via which they affect physiology and pathology. *Mus musculus* possesses two functional *Period* genes (*Per1* and *Per2*) whose double knockout (KO) leads to immediate and complete arrhythmicity, whereas single *Per1* or *Per2* KO animals display partial rhythmicity, including altered behavioral periods, delayed arrhythmicity, or residual ultradian behavioral patterns^[10–13]^. Similarly, the double KO of both *Cry1* and *Cry2* is required for complete arrhythmicity^[14,15]^, since single *Cry1* or *Cry2* KO only affects period length in an opposing manner while preserving oscillation^[15,16]^. *Bmal1* is an essential core clock component whose single KO causes immediate and complete arrhythmicity^[17]^. Although multiple animal models have consistently displayed arrhythmicity or altered rhythmicity, the effects of various KOs beyond the circadian rhythm manifest differently. For instance, muscular and skeletal degeneration are only observed following *Bmal1* KO^[17]^. Similarly, *Per* and *Bmal1* KO affect mood regulation, with *Per1,2*^-/-^ mice displaying increased anxiety without hyperactivity^[18]^ and shRNA-mediated *Bmal1* knockdown in the SCN increasing both anxiety and depression^[19]^. Furthermore, depending on testing time *Clock* delta19 mutants have been shown to display low anxiety and depression^[20]^, while *Rev-erba* KO mice were found to decrease anxiety and depression^[21]^. These conflicting findings highlight the need for an efficient and accessible strategy that can target multiple components of the molecular negative feedback loop to investigate local clocks.

The CRISPR-Cas9 system is derived from the adaptive immune system of prokaryotes and cleaves specific sequences in foreign DNA^[22]^. For genome editing, guide RNA (gRNA) containing a 20 bp spacer sequence followed by PAM 5’-NGG enables the Cas9-gRNA complex to recognize and be recruited to genomic loci complementary to the spacer sequence. Cas9 then generates a double strand break (DSB) that is corrected by a DNA repair system such as the non-homologous end joining pathway, which can generate a quasirandom indel at the DSB to potentially produce a frame-shift mutation^[22]^. In circadian genes, several applications of the CRISPR-Cas9 system has been reported to KO molecular clock components in a robust and efficient manner. Korge *et al*. first utilized CRISPR-Cas9-mediated gene targeting in chronobiology to generate *FBXL3* KO U2-OS cells^[23]^ and have recently extended this application to generate *CRY1* and *CRY2* KO U2-OS cells. The CRISPR-Cas9 system has also been used to generate KO animals lacking core clock genes in mice, macaques, monarch butterflies, and *Neurospora crassa*^[24–47]^. Virally delivered single guide RNAs (sgRNAs) targeting core clock components have also proven to be efficient *in vitro* and *in vivo*^[28,29]^; however, CRISPR-Cas9 system has only been applied to chronobiology for single genes at a time, thus limiting its ability to abolish the molecular clock as many of its components display genetic complementation.

In this study, we developed a CRISPR-Cas9-based single adeno-associated viral (AAV) system targeting the circadian clock (CSAC) using CRISPR-Cas9-mediated genome editing techniques and an AAV vector system to simultaneously KO multiple components of core clock machinery. We assembled a set of individual sgRNAs that can efficiently induce genomic mutation and confirmed reduced protein levels in the targeted genes. In addition, we demonstrated that CSAC can efficiently abolish the molecular and behavioral oscillation of the circadian rhythm *ex vivo* and *in vivo*, respectively. Thus, CSAC has the potential to improve our understanding of the effects of local clocks on physiology and pathology.

## RESULTS

### Design and evaluation of sgRNAs targeting core clock gene family members

To obtain a set of single guide RNAs (sgRNAs) capable of knocking out essential molecular components of the circadian clock, we designed sgRNAs targeting core clock genes (*Cry1, Cry2, Per1, Per2*, and *Bmal1*) using CHOPCHOP, a web-implemented target-site selecting algorithm for CRISPR-Cas9^[30]^. Among the predicted sgRNAs, we selected three targeting early exons to maximize the effect of the frame shift-mediated non-sense mutation (Fig. 1a). The individual sgRNAs were then cloned into a U6 promoter-driven sgRNA expression vector to evaluate their mutation-inducing efficiency (Fig. 1b). Each sgRNA-expressing plasmid was transfected into the Neuro2a neuroblastoma cell line with *SpCas9*-expressing plasmids. The mutation rate of the harvested genomic DNA was evaluated using Surveyor nuclease mismatch cleavage assay that can detect possible mutations induced in the genomic DNA of transfected cells, whether it be a single base conversion or an indel.

**Figure 1.**
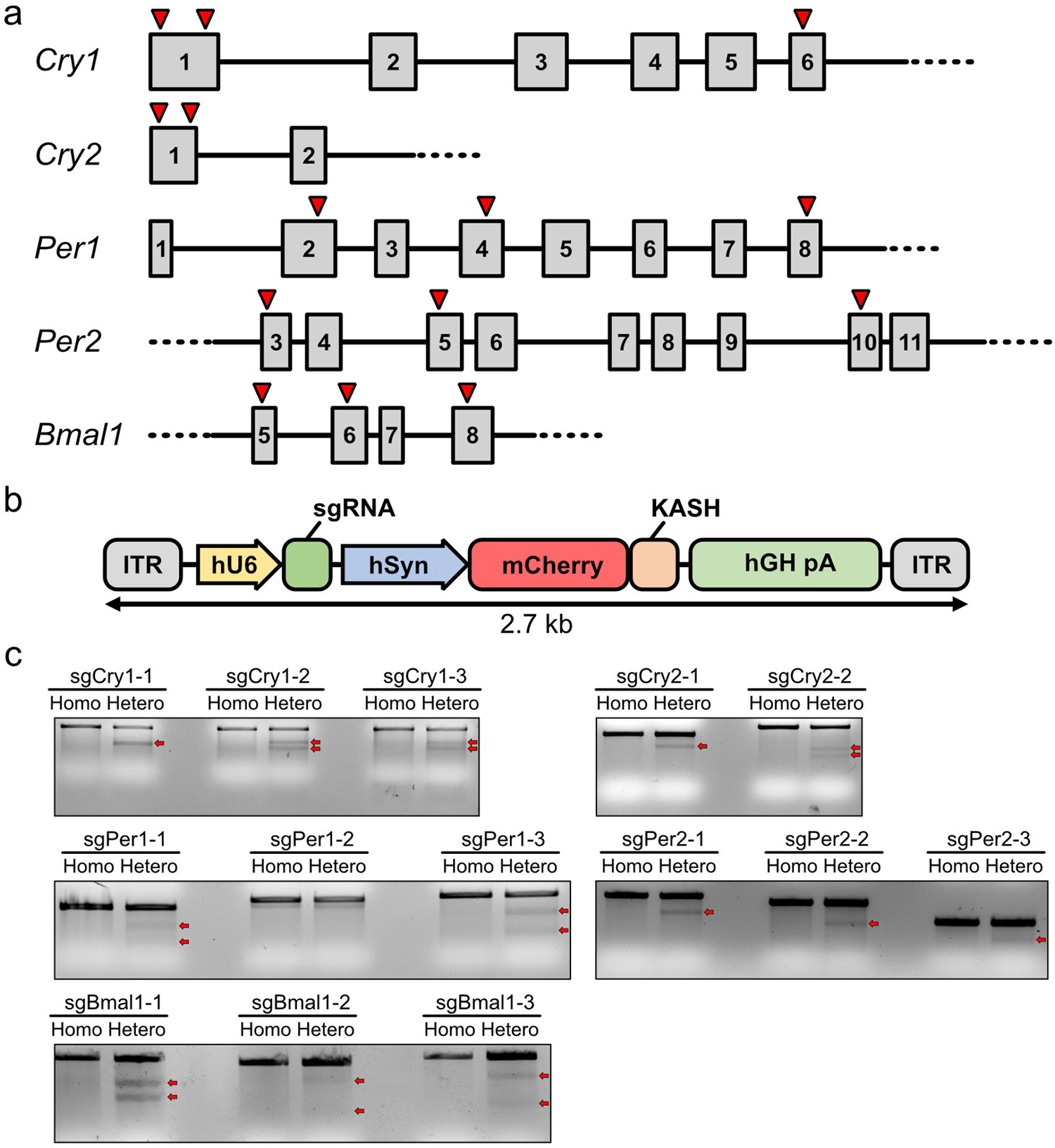
Design and evaluation of single guide RNAs (sgRNAs) targeting core molecular clock family members. **(a)** Genomic location of sgRNAs targeting *Cry1, Cry2, Per1, Per2*, and *Bmal1*. Gray boxes indicate exons. Lines indicate introns. Red arrows indicate the location of sgRNAs. **(b)** Structure of sgRNA expression vectors driven by the U6 promoter (hU6). ITR, inverted terminal repeat; hSyn, human synapsin promoter; KASH, Klarsicht ANC-1 syne homology domain for nuclear membrane targeting; hGH pA, poly adenylation signal derived from human growth hormone. **(c)** Mutation efficiency determined by Surveyor cleavage assays. Red arrows indicate cleaved PCR products, indicative of introduced mutations. Homo, homoduplex of PCR-amplified DNA fragments from the genomic DNA of non-transfected Neuro2a cells; Hetero, heteroduplex of PCR-amplified DNA fragments from the genomic DNA of non-transfected and sgRNA-expressing-plasmid-transfected Neuro2a cells.

All the genomic DNA of Neuro2a cells transfected with plasmids expressing sgRNA targeting *Cry1* (sgCry1) yielded cleaved DNA bands in the mutation mismatch cleavage assay (Fig. 1c, arrows). In the sgCry2 group, both sgRNAs caused cleavage, but not as abundantly as in the sgCry1 groups. Since a single AAV can harbor at most three sgRNA-expressing cassettes, we combined one sgRNA targeting *Cry1* with strong mutation-inducing efficiency (sgCry1-2) and two sgRNA targeting *Cry2* with modest mutation-inducing efficiency (sgCry2-1 and sgCry2-2) to increase the possibility of simultaneously knocking out both *Cry1* and *Cry2* (Fig. 1c). sgRNAs for *Pers* were selected in a similar manner. Since the sgPer1-3, sgPer2-1, and sgPer2-2 groups all produced intense cleavage bands, these three sgRNA targeting *Per*s were selected for the multiplexed AAV vector. For *Bmal1*, we combined two well-functioning sgRNAs targeting different *Bmal1* sites to increase the efficacy of *Bmal1* KO. The sgRNA sequences are shown in Table 1.

**Table 1.**
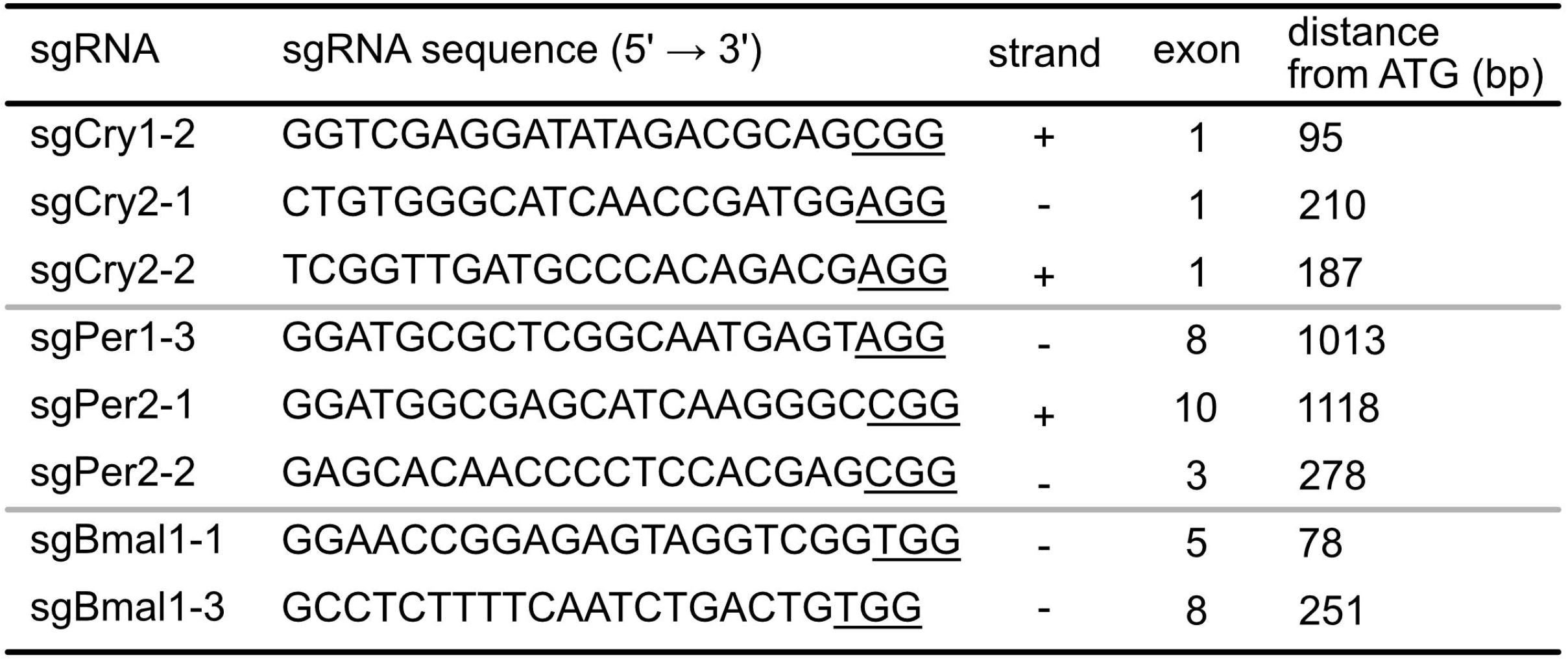
Nucleotide sequences of sgRNAs selected for CSAC construction. Underlined sequences indicated PAM sequence. Strand denotes whether sgRNA recognizes sense (+) or anti-sense (−) strand of target gene.

Next, we examined the genome editing efficacy of the multiplexed sgRNAs when reducing protein expression. Neuro2a cells stably expressing Cas9:EGFP (hereafter referred to as Neuro2a-Cas9 cells) were transfected with plasmids harboring multiplexed U6-mediated sgRNA expression cassettes (Fig. 2a) before protein extracts were isolated and subjected to western blot assays with antibodies against CRYs, PERs, or BMAL1. In Neuro2a-Cas9 cells transfected with plasmids expressing sgCrys, CRY1 and CRY2 protein levels were markedly lower than in cells transfected with the sgLacZ control (Fig. 2b). Similarly, the abundance of PER1 and PER2 protein in the sgPers-transfected group was less than half of that in the sgLacZ control group, while the sgBmal1s decreased BMAL1 levels. Consistently, mutation assays of genomic DNA from transfected cells and western blot assays indicated successful targeted gene KO; therefore, we named the multiplexed sgRNA-expressing AAV vector system CSAC.

**Figure 2.**
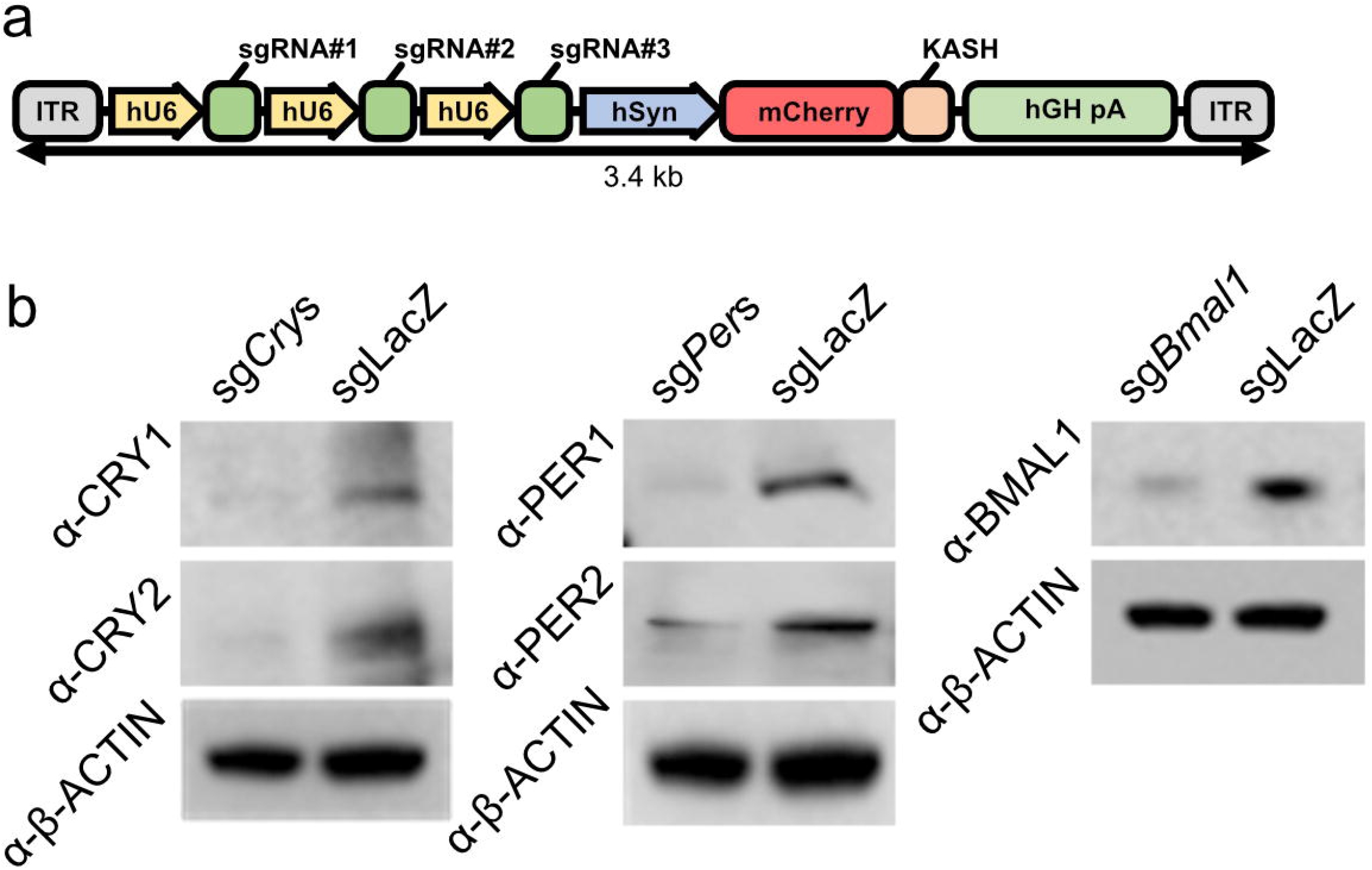
Reduced protein levels of core clock components by genome editing with a multiplexed sgRNA-expressing cassette. **(a)** Structure of multiplexed U6 promoter-driven sgRNA expression cassettes for AAV packaging. **(b)** Representative western blots of protein extracts from Neuro2a-Cas9 cells transfected with CSAC-*Crys* (left), CSAC-*Pers* (middle), or *CSAC-Bmal1* (right). Proteins from CSAC-*Crys*-infected cells were probed with anti-CRY1 and −CRY2 antibodies. Proteins from CSAC-*Pers*-infected cells were probed with anti-PER1 and −PER2 antibodies. Proteins from CSAC-*Bmal1*-infected cells were probed with anti-Bmal1 antibodies, with anti-β-actin antibodies used as an internal control for total protein abundance.

### CSAC dampened molecular clock oscillation in SCN slice cultures

To determine whether the depleted protein levels caused by CSAC could abolish central molecular clock oscillation at a population level, we monitored the oscillation of bioluminescence emitted from organotyptic SCN slice cultures from neonatal knock-in animal expressing a PER2 and LUCIFERASE fusion protein and constitutively expressing Cas9 (PER2::LUC;Cas9; Fig. 3a)^[31,32]^. The bath application of CSAC followed by a 2 week incubation enabled the widespread expression of fluorescent reporter proteins, indicating sgRNA expression in the SCN (Fig. 3b). The SCN explants expressing control sgLacZ exhibited robust circadian oscillation in PER2 expression during the recording session (Fig. 3c); however, PER2 oscillation was severely dampened in SCN cultures infected with either CSAC-*Cry*s or *CSAC-Bmal1* after the first peak induced by medium change, without a noticeable increase in PER2::LUC expression (Fig. 3d, e). To rule out the possibility that CSAC damages SCN cells or interferes with the genes regulating PER2 expression, we pharmacologically activated PER2 expression by administering forskolin, an adenylyl cyclase activator^[33]^. In all groups, forskolin treatment robustly induced PER2::LUC expression, regardless of CSAC. The statistical analysis of free-running PER2::LUC oscillation further clarified that CSAC induced dampening of the molecular clock, with CSAC-*Crys* and CSAC-*Bmal1* both significantly suppressing the amplitude of PER2::LUC oscillation (Fig. 3f). CSAC also affected the robustness of bioluminescence oscillation, although to a lesser degree than its impact on amplitude due to the succession of suppressed peaks (Fig. 3g), and did not significantly affect the period of residual molecular oscillation (Fig. 3h). Together, these findings indicate that CSAC effectively and specifically dampens molecular clock oscillation *ex vivo*.

**Figure 3.**
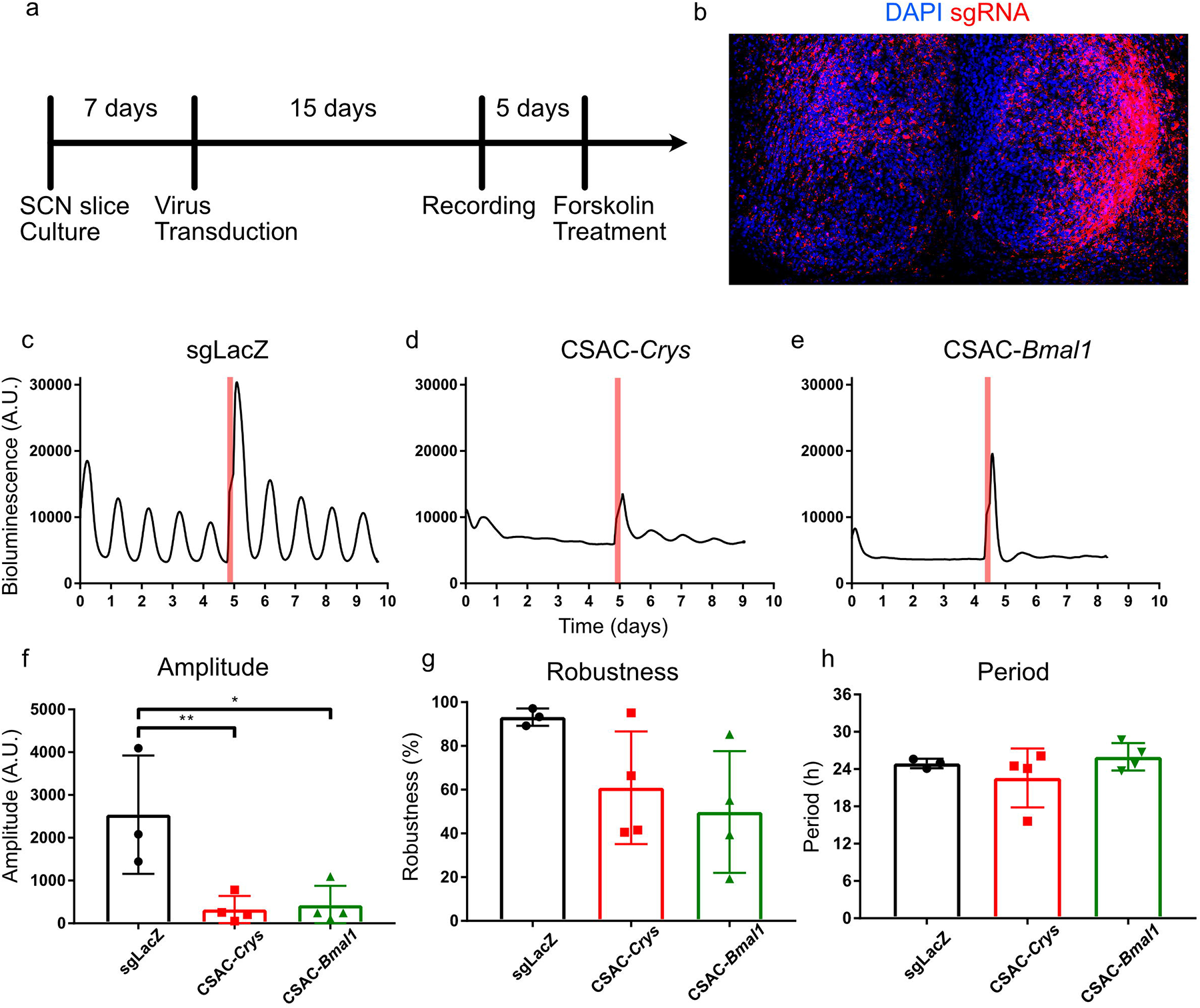
CSAC abolishes molecular clock oscillation *ex vivo*. **(a)** Timeline of organotypic slice culture preparation and viral infection. **(b)** Representative confocal microscopy image of organotypically cultured SCN from mPER2::LUC;Cas9 mice. **(c-e)** Representative profiles of PER2::LUC mice with CSAC. Bioluminescence signals from SCN explant cultures infected with either sgLacZ (**c**), CSAC-*Crys* (**d**), or *CSAC-Bmal1* (**e**) were continuously monitored. Red lines indicate forskolin treatment. **(f-h)** Oscillation amplitude (**f**), robustness (**g**), and period (**h**) in PER2::LUC slice cultures infected with AAV analyzed using Cosinor. \**p* < 0.05; ** *p* < 0.01 by Tukey’s post-hoc test (*n* = 3-4 slices per group).

### SCN-injected CSAC induced arrhythmicity in locomotor activity and body temperature

To demonstrate the utility of CSAC *in vivo*, we injected either CSAC-*Cry*s, CSAC-*Per*s, *CSAC-Bmal1*, or control AAV into the SCN of mice constitutively expressing Cas9 (Fig. 4a). AAV delivery was histologically verified by confocal microscopy of the brain sections containing the SCN (Fig. 4b). For the following analysis, only mice with viral infection sites covering the SCN were included (Fig. 4c-f).

**Figure 4.**
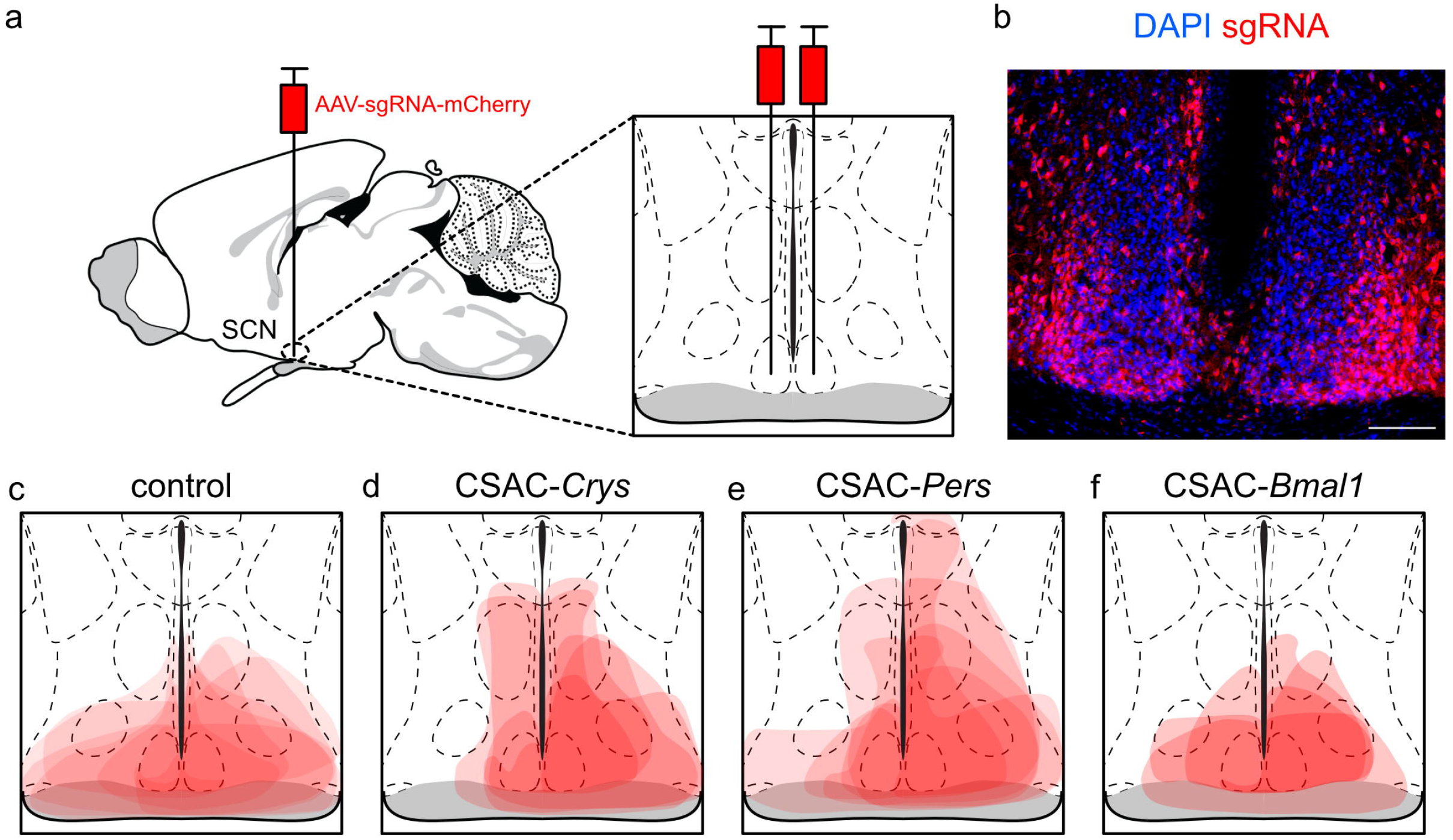
SCN injection with CSAC. **(a)** Schematic diagram of stereotaxic CSAC injection into the SCN. AAVs expressing sgRNAs targeting core clock components and the mCherry fluorescent marker were bilaterally injected into the SCN of Cas9-expressing mice. **(b)** Representative confocal image of the brain section containing CSAC-injected SCN. sgRNA-expressing cells are co-labeled with mCherry as a surrogate marker. Scale bar = 100 μm. **(c-f)** Injection site of control virus (**c**), CSAC-*Crys* (**d**), CSAC-*Pers* (**e**), and *CSAC-Bmal1* (**f**). Light red indicates the infected areas of each animal.

To determine the effect of CSAC on the behavioral and physiological aspects of circadian rhythm in awake behaving animals, we used a telemetry-based method to simultaneously record locomotor activity and body temperature. Circadian locomotor activity was normal in control animals injected with AAV expressing fluorescent protein (mCherry; Fig. 5a), which exhibited light-suppressed locomotion during light phase and a distinct behavioral onset that aligned well with the point at which the light was turned off. Under constant darkness, the control animals clearly displayed a circadian wake and resting cycle with a period slightly shorter than 24 h, characteristic of the C57BL/6J strain^[34]^. However, the mice injected with CSAC-*Cry*s or CSAC-*Per*s displayed arrhythmicity under constant darkness (DD) (Fig. 5b, c). These mice were subjected to a 12 h light/dark (LD) cycle to maintain a daily pattern of locomotor activity, which was completely abolished under DD. In the CSAC-*Bmal1-*injected group, circadian locomotor activity largely disappeared during DD and generally displayed lower locomotor activity during LD than the other groups (Fig. 5d). Chi-square periodogram analyses clearly indicated that CSAC disrupted circadian locomotor activity^[35]^. In the control animals, the average Qp value significantly peaked slightly before 24 h (Fig. 5e), typical of normal circadian locomotor activity, whereas the average Qp values of CSAC-*Crys*, CSAC-*Pers*, and *CSAC-Bmal1* groups were not significant at any time point (Fig. 5f-h).

**Figure 5.**
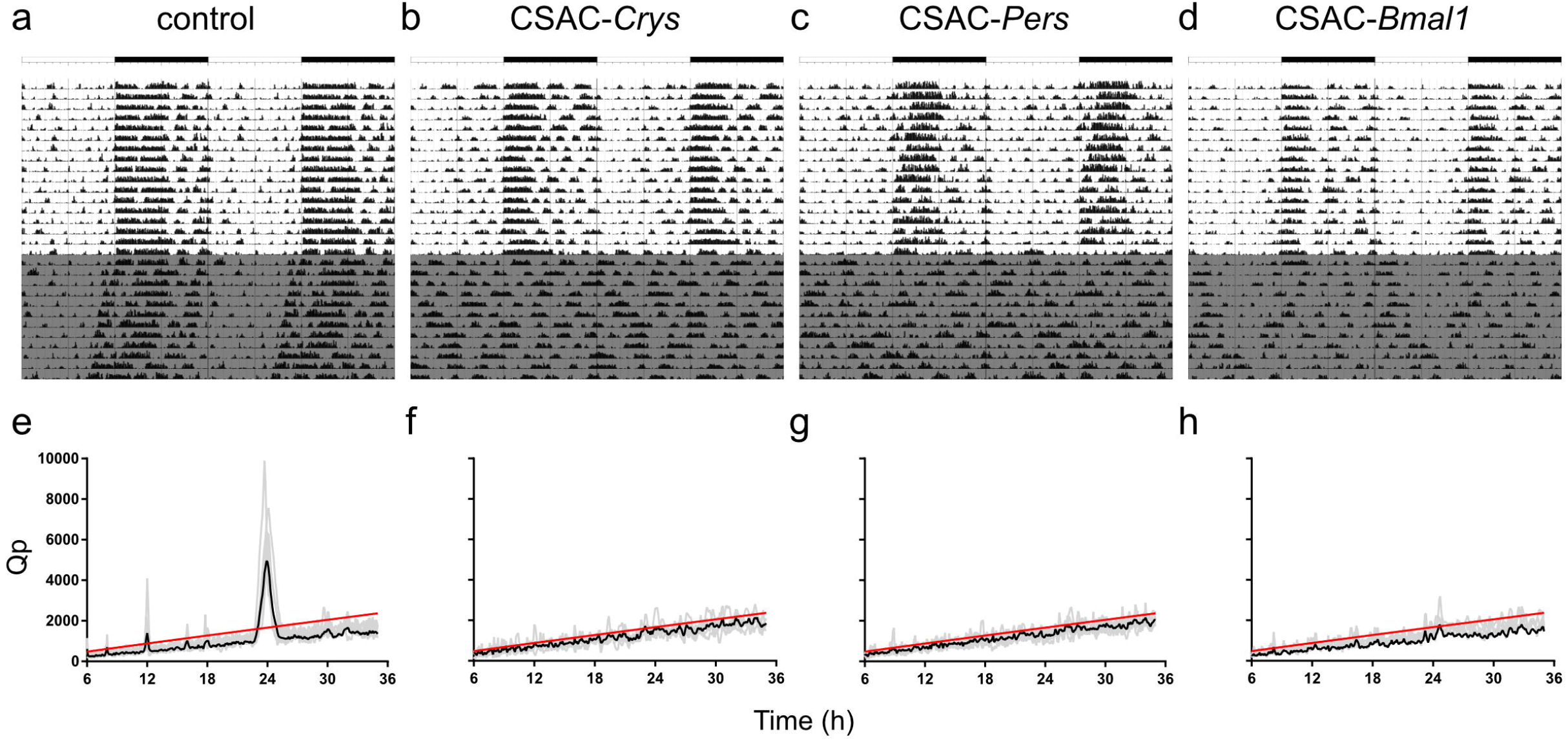
Circadian locomotor activity in SCN-targeted mice. **(a-d)** Representative locomotor pattern shown as a double plot of activity over 28 days in control (**a**), CSAC-*Crys*- (**b**), CSAC-*Pers*- (**c**), and *CSAC-Bmal1-* (**d**) injected mice. Gray indicates periods of constant darkness. **(e-h)** Chi-square periodogram of locomotor activity during constant darkness in control (**e**), CSAC-*Crys*- (**f**), CSAC-*Pers*- (**g**), and *CSAC-Bmal1-* (**h**) injected mice. Gray indicates the periodogram of an individual mouse. Black line indicates the averaged spectrogram of each group. Red line indicates the statistical significance threshold (*n* = 4-8 mice per group).

Body temperature largely followed the same pattern as locomotor activity in the experimental animals. For instance, the control animals displayed a typical circadian actogram under both LD and DD conditions (Fig. 6a), whereas the CSAC-*Cry*s, CSAC-*Per*s, and *CSAC-Bmal1* groups lacked a circadian change in body temperature under free-running conditions without a light cue (Fig. 6b-d). Again, Chi-square periodograms supported these findings, with the average Qp value of the control animals exceeding the significance threshold at approximately 24 h, unlike the mice in the CSAC-*Crys*, CSAC-*Pers*, or CSAC-*Bmal1* groups (Fig. 6e-h).

**Figure 6.**
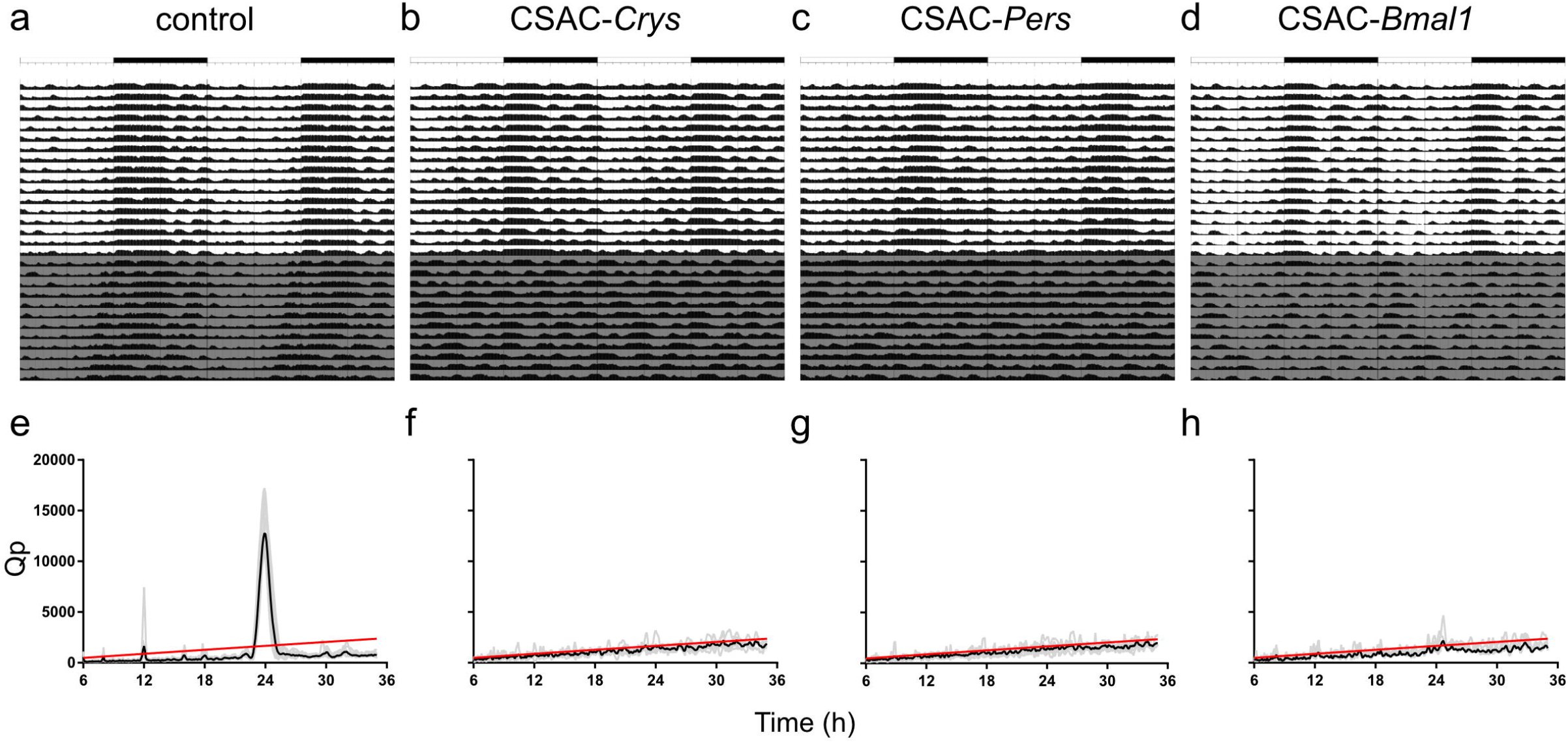
Circadian body temperature of SCN-targeted mice. **(a-d)** Representative body temperature profile shown as a double plot over 28 days for control (**a**), CSAC-*Crys*- (**b**), CSAC-*Pers*- (**c**), and *CSAC-Bmal1-* (**d**) injected mice. Gray indicates periods of constant darkness. **(e-h)** Chi-square periodogram of body temperature during constant darkness in control (**e**), CSAC-*Crys*- (**f**), CSAC-*Pers*- (**g**), and *CSAC-Bmal1-* (**h**) injected mice. Gray line indicates the periodogram of an individual mouse. Black line indicates the averaged spectrogram of each group. Red line indicates the statistical significance threshold (*n* = 4-8 mice per group).

To evaluate whether CSAC-mediated depletion of circadian proteins in the SCN affected baseline activity levels or temperature, we calculated average activity (Fig. 7a, c) and body temperature (Fig. 7b, d), respectively. We found that average locomotor activity and body temperature did not significantly differ between the control and CSAC groups. Together, these findings indicate that CSAC injection into the SCN efficiently and specifically suppressed the physiological hallmarks of circadian rhythm *in vivo* without affecting average locomotor activity or body temperature.

**Figure 7.**
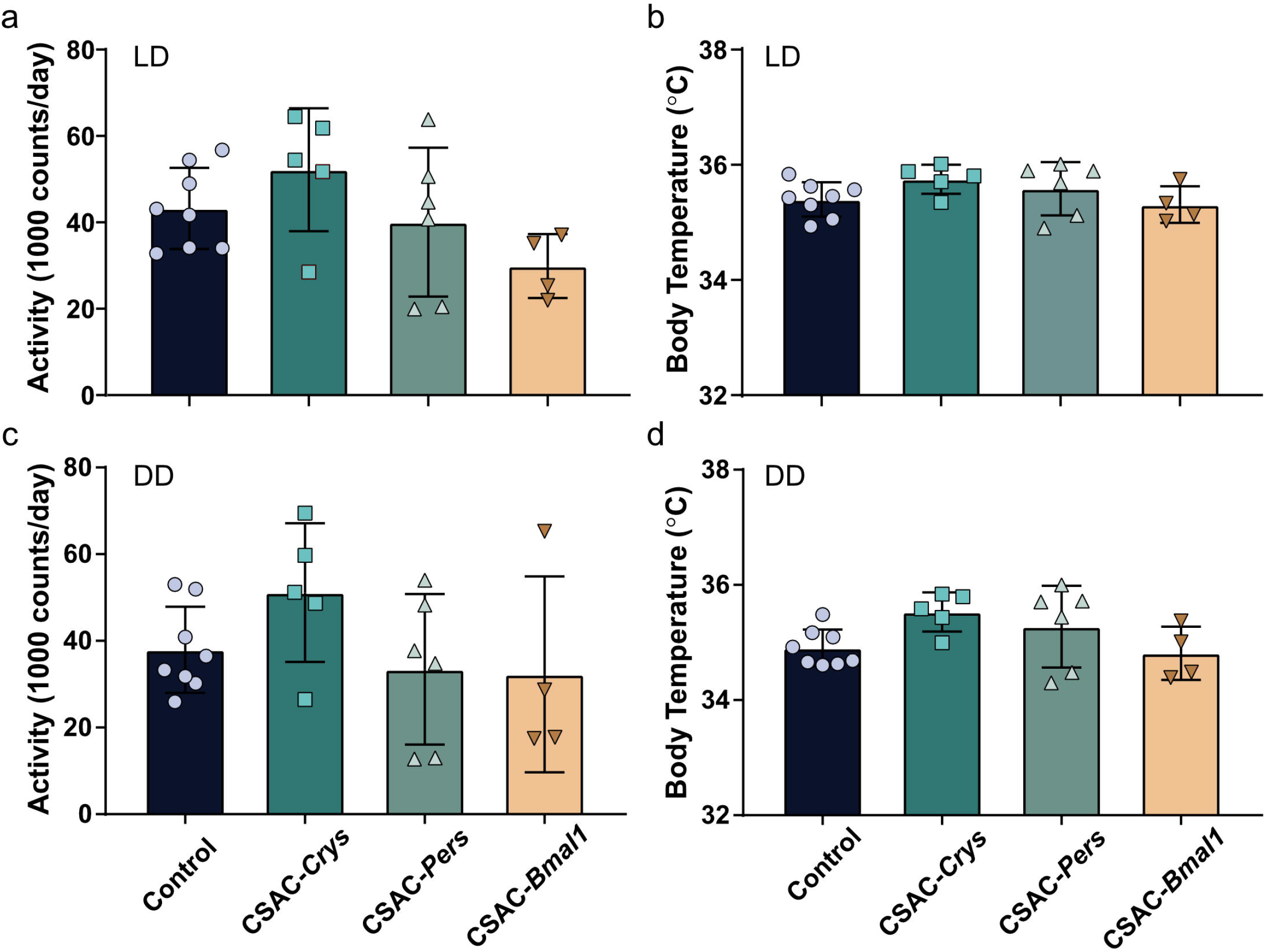
Average locomotor activity and body temperature. **(a)** Average locomotor activity under a 12h light/dark (LD) cycle. **(b)** Average body temperature under an LD cycle. **(c)** Average locomotor activity under constant darkness (DD). **(d)** Average body temperature under DD. Error bar = standard deviation (*n* = 4-8 mice per group).

## DISCUSSION

In this study, we devised CSAC, a system of multiplexed sgRNAs targeting core clock components packaged in a single AAV, to abolish molecular clock oscillation by inducing null mutations into several core clock genes. We selected the multiplexed sgRNAs based on their mutation-introducing efficacy and target loci to maximize the possibility of null mutations in the target genes. By targeting multiple core clock genes (*Cry*s, *Pers*, and *Bmal1*), we decreased the expression levels of the targeted genes *in vitro* and *in vivo*. In addition, we demonstrated the value of CSAC in multiple systems including cell lines, organotypic tissue cultures, and live animals.

To our knowledge, we are the first to develop a set of *in vivo* applicable viral vectors targeting members of core clock gene families employing CRISPR-Cas9-based genome editing, while CRISPR-Cas9-mediated genome editing has already been applied to *in vitro* system or *in vivo* single targeting in the field of chronobiology. Korge *et al*. first reported the CRISPR-Cas9-based genomic alteration of the circadian rhythm by targeting F-box and leucine-rich repeat protein 3 (*Fbxl3*)^[23]^. To do so, they transfected U2-OS cells stably expressing a Bmal1-Luciferase reporter with a lentivirus expressing Cas9 and two sgRNA targeting different sites in the *Fbxl3* gene, and found that CRISPR-Cas9-based *Fbxl3* KO dampened *Bmal1* promoter-driven luminescence oscillation. The same group then utilized a CRISPR-Cas9 system to generate CRY1 and CRY2 double KO cells^[36]^. Targeting *Clock* has also demonstrated efficacy in mouse embryonic stem cells when investigating the non-clock functions of the *Clock* gene^[25]^, while Lee and colleagues expanded the repertoire of direct CRISPR-Cas9 targets by designing an adenoviral vector system for efficiently generating null mutant cell lines^[28]^. Similarly, Herzog and colleagues used AAV vectors to target *Bmal1* in astrocyte-specific *Cre*-driven Cas9-expressing mice^[29]^, allowing genome editing to be applied *in vivo* in adult animals. These studies provided a foundation for the CSAC developed here, which offers an advanced approach for knocking out the molecular clock *in vivo*.

The efficient KO of core clock components by CSAC in Cas9-expressing mice indicates that the multiplexed sgRNAs can properly recognize target genomic loci. The expanding repertoire of CRISPR-Cas9 technologies suggest that CSAC could be utilized for a variety of applications beyond the KO of target genes^[37]^. In this study, we used *Streptococcus pyrogen* Cas9 (SpCas9), which contains two nuclease domains, RuvC and HNH. Mutations in these two domains (Cas9 D10A H840A) can lead to a catalytically dead Cas9 mutant (dCas9) that can bind to target genomic loci as dictated by sgRNA, but cannot generate a DSB^[38]^. The conjugation of dCas9 to either a transcriptional activator (dCas9-VP64 and dCas9-p65) or repressor (dCas9-KRAB) has been reported to efficiently induce or suppress specific sgRNA-target genes^[39–41]^; thus, intra-coding sequence-targeting sgRNAs such as CSAC could be used to regulate gene expression. When combined with the conditional expression of dCas9 derivatives using chemical or optogenetic approaches^[37]^, CSAC may be able to temporally control the expression of core clock genes to mimic the circadian regulation of gene expression. In addition, Cas9 variants targeting RNA could be utilized to visually track *Pers* or *Crys* mRNA in live cells^[42]^. Considering the ever-growing molecular repertoire of CRISPR-Cas9 technologies, we believe that the applications of CSAC will expand accordingly.

In this study we designed and validated CSAC as a powerful and accessible tool for ablating the molecular clock. We demonstrated that CSAC can efficiently reduce the protein expression of core clock genes, dampen their molecular oscillation *ex vivo*, and disrupt physiological circadian outputs, such as locomotor activity and body temperature. Combined with a Cre-dependent Cas9 expression system and Cre driver line, CSAC offers a potent and practical tool for studying the role of the molecular clock in specific cell types of interest and its applications will expand alongside advances in CRISPR-Cas9-based genome editing.

## METHODS

### Animals

All mice were born and reared in standard mouse cages (16 × 36 × 12.5 cm) with food and water available *ad libitum*. Mice were weaned at 3-4 weeks of age and housed together with same sex siblings with up to four animals per cage. Mice were maintained under a 12 h light/dark cycle at 22 ± 1 □. All procedures were approved by the Institutional Animal Care and Use Committee of Daegu Gyeongbuk Institute of Science and Technology (DGIST). PER2::LUC knock-in mice were a generous gift from Joseph Takahashi ^[31]^. Cas9-expressing mice were provided by Feng Zhang ^[32]^.

### Cell culture and mutation detection

Cells were cultured as described previously, with minor modifications^[43]^. Neuro2a and Neuro2a Cas9-GFP-Hygor^res^(CSC-RO0033, Genecopeoia, Rockville, MD, USA) cells were cultured in Dulbecco’s modified Eagle medium (DMEM) (Hyclone, Chicago, IL, USA) containing 1× penicillin/streptomycin (Capricorn Scientific, Ebsdorfergrund, Germany), 10 mg/mL hygromycin (H-34274, Merck Millipore, Burlington, MA, USA), and 10 % fetal bovine serum (FBS) (Hyclone) at 37 □ in 5 % CO_2_. Neuro2a-Cas9 cells were transfected with pAAV-U6-sgBMAL-hSyn-mCherry, pAAV-U6-sgCRY-hSyn-mCherry, pAAV-U6-sgPER-hSyn-mCherry, or pAAV-U6-sgLacZ-hSyn-mCherry using Lipofectamine 2000 transfection reagent (#11668019, Thermo Scientific, Waltham, MA, USA), according to the manufacturer’s recommendations. Cells were collected after 48 h and genomic DNA was prepared using a tissue mini kit according to the manufacturer’s instructions (Cosmo Genetech, Seoul, South Korea). Surveyor mutation detection assays were performed according to the manufacturer’s instructions (Integrated DNA Technologies, Coralville, IA, USA).

### Plasmids and AAV production

DNA oligomers of sgRNAs designed using CHOPCHOP^[30]^ with adaptors for cloning were commercially synthesized by Bionics (Seoul, South Korea). After annealing, the DNA fragments containing sgRNAs were cloned into the sgRNA expression vector, pAAV-hU6-gRNA-hSyn-mCherry-KASH. The U6-sgRNA expression cassettes were multiplexed using the Goldengate assembly method, yielding three multiplexed cassettes assembled in pAAV-(hU6-gRNA)X3-hSyn-mCherry-KASH. The plasmid DNA sequences were validated by Sanger sequencing (Macrogen, Seoul, South Korea). AAVs were produced using the triple transfection method and purified using the iodixanol gradient method^[44]^. All AAV vectors administered in the paper was tittered between 10^12^ and 10^13^ viral genome copies per milliliter (GC/mL), as quantified by qPCR: AAV-DJ-hU6-sgLacZ-hSyn-mCherry-KASH 1.8 × 10^13^ GC/mL; AAV-DJ-CSAC-Crys-hSyn-mCherry-KASH 1.1 × 10^13^ GC/mL; AAV-DJ-CSAC-Pers-hSyn-mCherry-KASH 3.1 × 10^13^ GC/mL; AAV-DJ-CSAC-sgBmal1-hSyn-mCherry-KASH 6.6 × 10^12^ GC/mL; AAV-2-hSyn-mCherry 4.7 × 10^12^ GC/mL.

### Western blot analysis

Cell were washed twice with phosphate-buffered saline and lysed with sonication in radioimmunoprecipitation assay (RIPA) buffer (#9806, Cell Signaling, Danvers, MA, USA), before being incubated on ice for 30 min. Lysates were centrifuged to obtain the supernatants, which were boiled for 5 min in Laemmli sample buffer (#NP0007, Invitrogen, Waltham, MA, USA). Protein were separated on 4-15 % Tris-glycine (TG) gels (#45101080003-2, SMOBIO, Hsinchu, Taiwan) and electro-transferred onto polyvinylidene fluoride (PVDF) membranes (#A16646282, GE Healthcare, Chicago, IL, USA) in TG buffer. The membranes were blocked in Tris buffered with Tween-20 (TTBS) with 1 % bovine serum albumin (BSA) and incubated overnight at 4 □ in TTBS with 0.1% BSA and the following primary antibodies: mPer1 (#AB2201, Merck Millipore), PER2 (#AB2202, Merck Millipore), BMAL1 (#NB100-2288, Novus, Centennial, CO, USA), CRY1 (#AF3764-SP, R&D Systems, Minneapolis, MN, USA), CRY2 (#HPA037577, Atlas Antibodies, Bromma, Sweden), and β-actin-HRP (#SC-47778, Santa Cruz, Dallas, TX, USA). Anti-Rabbit (SKU#31460, Thermo Scientific) or antigoat (Jackson Laboratory, Bar Harbor, ME, USA) secondary antibodies were used. Protein bands were detected using enhanced chemiluminescence (ECL) Select Solution (GE healthcare) and an ECL Pico System (ECL-PS100, Dongin, Seoul, Korea).

### Bioluminescence monitoring of organotypic SCN slice cultures

SCN explant cultures were prepared and monitored similarly to a previous report with minor modifications^[45]^. One-week-old mPer2::LUC;Cas9 mice were sacrificed and their brains were quickly removed before being chilled on ice and moistened in Gey’s Balanced Salt Solution (GBSS), supplemented with 0.01 M HEPES and 36 mM D-glucose, and aerated with 5 % CO2 and 95 % O2. The brains were then coronally cut into 400-μm thick slices using a Leica VT1000 S vibratome (Leica, Wetzlar, Germany). The slices were maintained on a culture insert membrane (Millicell-CM; Millipore, Bedford, MA, USA) and dipped into culture medium (50 % minimum essential medium, 25 % GBSS, 25 % horse serum, 36 mM glucose, and 1 × antibiotic-antimycotic) at 37 °C. The SCN slices were cultivated for more than two weeks before being used in experiments. To quantitatively measure bioluminescence, the SCN cultures were maintained in a 35-mm petri dish with 1 mL culture medium containing 0.3 mM D-luciferin (Promega, Madison, WI, USA) at 36 °C. Light emission was measured and integrated for 1 min at 10-min intervals using a dish-type wheeled luminometer (AB-2550 Kronos-Dio; ATTO, Tokyo, Japan). Unless otherwise specified, all culture media and supplements for organotypic slice culture were purchased from Thermo-Fisher Scientific.

SCN slices were virally transduced after 7 days of stabilization, as described previously^[46]^. AAVs were dropped directly onto the surface of each SCN slice (1-2 μL per slice) in a 35-mm dish and incubated for 15 days prior to bioluminescence monitoring.

### Stereotaxic injection

Surgery was carried out using aseptic techniques. Briefly, mice were anesthetized with a mixture of ketamine (100 mg/kg) and xylazine (10 mg/kg) and placed in a stereotaxic apparatus (RWD Life Science, Shenzhen, China) where their brains were injected with AAVs (bilateral injection, 250 nL each) using Nanoliter injector system (WPI, Sarasota, FL, USA) at a rate of < 0.1 μL/min in the SCN (AP = −0.8 mm, ML = ±0.22 mm from bregma, DV = - 5.85 mm from brain surface).

### Measurement of locomotor activity and body temperature

The locomotor activity and body temperature of the mice were measured using E-mitter, a radio transmitter-based telemetry system (Starr Life Science, Oakmont, PA, USA). E-mitter was implanted beneath the skin on the backs of the mice using aseptic techniques under general anesthesia induced by intraperitoneal ketamine (100 mg/kg) and xylazine (10 mg/kg) injection^[47]^. After implantation, the mice recovered for at least one week and acclimatized in a regular 12 h light/dark cycle. Activity and temperature data detected by the implanted sensor were transmitted to a receiver (ER-4000 engergizer receiver, Starr Life Science). Data acquisition and digital transformation was performed using VitalView software every 6 min (Starr Life Science).

### Tissue sample preparation and confocal microscopy

After three weeks to allow viral expression, mice were intracardially perfused with 4 % paraformaldehyde and post-fixed at 4 □ overnight. Coronal sections (50-μm thick) were obtained using a Leica VT1000 S vibratome (Leica, Wetzlar, Germany). The nucleus was counterstained with 4’6’-diamidino-2-phnylindole (DAPI) and images were taken using a Nikon C2^+^ confocal microscope system and linearly adjusted with Fiji^[48]^.

### Data and statistical analysis

Chi-square periodogram analysis was performed using the xsp package^[49]^ in R^[50]^. Real-time bioluminescence was analyzed by the cosinor procedure^[51, 52]^. Graphs were constructed using Prism 8 (GraphPad, SanDiego, CA, USA) or ggplot2^[53]^. The threshold for statistical significance was set at *P* < 0.05.

## ACKNOWLEDGMENTS

This work was supported by the Korean National Research Foundation (NRF-2018M3C7A1022310 and NRF-2019M3C1B8090845), intramural grants from DGIST (20-RT-01 and 20-CoE-BT-03), and the KBRI basic research program (17-BR-04). We thank Kyojin Ku for technical assistance in preparation of SCN organotypic cultures.

## AUTHOR CONTRIBUTIONS

B.K. designed the work, performed the experiments, and analyzed the data. J.K, M.C., I.P., M.C. performed the experiments. K.K. conceived the work. H.K.C. conceived, designed, and drafted the work.

## ADDITIONAL INFORMATION

### Data availability

All data generated and analysed during this study are included in this article.

